# GP96 drives exacerbation of secondary bacterial pneumonia following influenza A virus infection

**DOI:** 10.1101/2020.09.14.297192

**Authors:** Tomoko Sumitomo, Masanobu Nakata, Satoshi Nagase, Yuki Takahara, Mariko Honda-Ogawa, Yasushi Mori, Masaya Yamaguchi, Shigefumi Okamoto, Shigetada Kawabata

## Abstract

Influenza A virus (IAV) infection predisposes the host to secondary bacterial pneumonia, known as a major cause of morbidity and mortality during influenza epidemics. Analysis of interactions between IAV-infected human epithelial cells and *Streptococcus pneumoniae* revealed that infected cells ectopically exhibited the endoplasmic reticulum chaperon GP96 on the surface. Importantly, efficient pneumococcal adherence to epithelial cells was imparted by interactions with extracellular GP96 and integrin α_V_, with the surface expression mediated by GP96 chaperone activity. Furthermore, abrogation of adherence was gained by chemical inhibition or genetic knockout of GP96, as well as addition of RGD peptide. Direct binding of extracellular GP96 and pneumococci was shown to be mediated by pneumococcal oligopeptide permease components. Additionally, IAV infection induced activation of calpains and Snail1, which are responsible for degradation and transcriptional repression of junctional proteins in the host, respectively, indicating increased bacterial translocation across the epithelial barrier. Notably, treatment of IAV-infected mice with the GP96 inhibitor enhanced pneumococcal clearance from lung tissues and ameliorated lung pathology. Taken together, the present findings indicate a viral-bacterial synergy in relation to disease progression and suggest a paradigm for developing novel therapeutic strategies tailored to inhibit pneumococcal colonization in an IAV-infected respiratory tract.

## Introduction

Secondary bacterial infection following influenza infection is associated with high rates of morbidity and mortality in elderly as well as chronically ill individuals. During the 1918 influenza pandemic, *Streptococcus pneumoniae* was identified as the predominant pathogen in more than 95% of all fatal cases^1^. Despite development of effective vaccines and potent antibacterial agents, during the 2009 H1N1 pandemic, bacterial pneumonia was a complication in one-quarter to one-half of severe and fatal cases^2^. The underlying mechanism of the viral-bacterial synergy leading to disease progression has remained elusive, thus hampering production of effective prophylactic and therapeutic intervention options.

Nasopharyngeal pneumococcal colonization is a major predisposing factor related to viral upper respiratory tract infections such as influenza^3, 4^. A preceding influenza virus infection can induce excess pneumococcal acquisition and carriage of the nasopharynx, which in turn promotes bacterial dissemination to the lungs. Although respiratory epithelium provides a physical barrier against most human pathogens, the influenza virus prefers to replicate in epithelial cells, leading to direct damage of airway epithelium^5, 6^. Virus-induced epithelial damage and exfoliation provide increased receptor availability for bacteria, resulting in establishment of bacterial colonization and onset of invasive diseases. For example, the influenza virus neuraminidase cleaves sialic acid glycoconjugates on airway epithelial cells as well as mucins, which facilitates not only bacterial adherence to cryptic receptors but also their proliferation in the upper respiratory tract^7^. Among bacterial receptors appearing on cell surfaces during influenza infection, platelet-activating factor receptor (PAFR) gained attention as a possible therapeutic target^8^. Extracellular PAFR binds to phosphorylcholine embedded in the cell walls of numerous respiratory bacterial pathogens, which subsequently accelerates lung bacterial burden and bacteremia, and increases mortality risk. However, previous studies have found that genetic knockout or pharmacological inhibition of PAFR had no effect on susceptibility of mice to secondary bacterial pneumonia, implicating multifaceted mechanisms related to the synergism between influenza viruses and bacterial pathogens^9, 10^.

A dual viral-bacterial infection causes dysfunction of the epithelial-endothelial barrier and, consequently, exudation of fluids, erythrocytes, and leukocytes into alveolar spaces, leading to gas exchange impairment and severe respiratory insufficiency. Indeed, pulmonary edema and hemorrhage are commonly found in autopsy examinations^1^. The physical barrier function of airway epithelium is provided by four types of cell-cell junctions; tight, adherens, and gap junctions, and desmosomes. Influenza virus-induced disruption of the pulmonary barrier is associated with a loss of integrity of claudin-4, a tight junctional protein^11^. Moreover, interaction between the PDZ-binding motif of the avian influenza virus NS1 protein and the PDZ domain present in tight junctional proteins has been shown to destabilize epithelial junctional integrity^12^. Although it is generally accepted that viral-induced epithelial cell damage allows for bacterial invasiveness, the molecular mechanisms involved in dysfunction of the alveolar epithelial barrier followed by bacterial dissemination remain largely unknown.

In the present study, novel findings showing that glycoprotein 96 (GP96), a host chaperone protein, is involved in exacerbation of bacterial pneumonia following influenza A virus (IAV) infection are presented. Interactions of pneumococci with extracellular GP96 and integrins, whose surface expressions are mediated by the chaperone activity of GP96, were found to promote pneumococcal adherence to IAV-infected epithelial cells. Also, inhibition of GP96 rendered IAV-infected cells as well as tested mice less susceptible to *S. pneumoniae* infection. Accordingly, GP96 is considered to be a potential target for therapeutic strategies for treating patients with superinfection. Furthermore, to the best of our knowledge, the present results are the first to demonstrate that viral infection induces calpain/Snail1-dependent dysfunction of the pulmonary epithelial barrier, thus providing a route for secondary bacterial translocation into deeper tissues via paracellular junctions. Together, these findings suggest an underlying mechanism responsible for polymicrobial synergy in cases of secondary bacterial pneumonia.

## Results

### Influenza A virus infection induces surface display of GP96 on alveolar epithelial cells

Host inflammatory response to a viral infection leads to increased or ectopic expressions of multiple proteins that serve as host receptors for bacteria. As a first step toward understanding the pathogenesis of bacterial pneumonia following influenza, we attempted to determine which proteins were exposed on the surface of alveolar epithelial cells following viral infection. Human A549 alveolar epithelial cells were infected with influenza A virus (IAV), followed by exposure to a membrane-impermeable biotinylation reagent. Cell surface proteins were then obtained using streptavidin beads, and subjected to SDS-PAGE and silver staining, which showed several different upregulated proteins on the surfaces of IAV-infected epithelial cells (Fig. 1a, arrows). Mass spectrometry analysis of these proteins revealed peptides corresponding to an endoplasmic reticulum protein, components of intermediate filaments, a glycolytic protein, and an oxidative stress-related protein. Among the host molecules, we focused on the human endoplasmic reticulum (ER) chaperon GP96, also referred to as GRP94 or endoplasmin, in further examinations. Although GP96 has been found to be mainly localized in the ER, abundant evidence presented indicates that it is also exposed on the surface of different cell types under particular conditions, such as infection, cell activation, and necrotic cell death^13^. Surface-displayed GP96 is frequently exploited as a receptor for bacterial pathogens, including *Listeria monocytogenes*^14^ and *Neisseria gonorrhoeae*^15^. To examine whether exposed GP96 serves as a receptor for *S. pneumoniae*, bacterial adherence to the apical surface of epithelial cells infected with or without IAV was examined. *S. pneumoniae* showed more efficient adherence to IAV-infected as compared to non-infected cells, while the enhanced bacterial association was reduced by pretreatment with either a pharmacological inhibitor of GP96 (Fig. 1b) or anti-GP96 antibody (Fig. 1c). On the other hand, a preceding IAV infection had no effect on the ability of *S. pneumoniae* to adhere to GP96 knockout cells (Fig. 1d), indicating it to be a critical factor for bacterial colonization on IAV-infected alveolar epithelial cells. To visualize GP96 distribution and pneumococci, non-permeabilized cells were stained with anti-GP96 and anti-pneumococcal capsule antibodies, respectively (Fig. 1e). GP96 was poorly visualized on the surface of non-infected cells, whereas its surface expression was markedly increased in response to IAV infection. Notably, co-localization of pneumococci with GP96 was observed in superinfected cells. Based on these results, we speculated that redistribution of GP96 on epithelial surfaces caused by IAV infection has a crucial role in secondary bacterial colonization in the lungs.

**Fig. 1.**
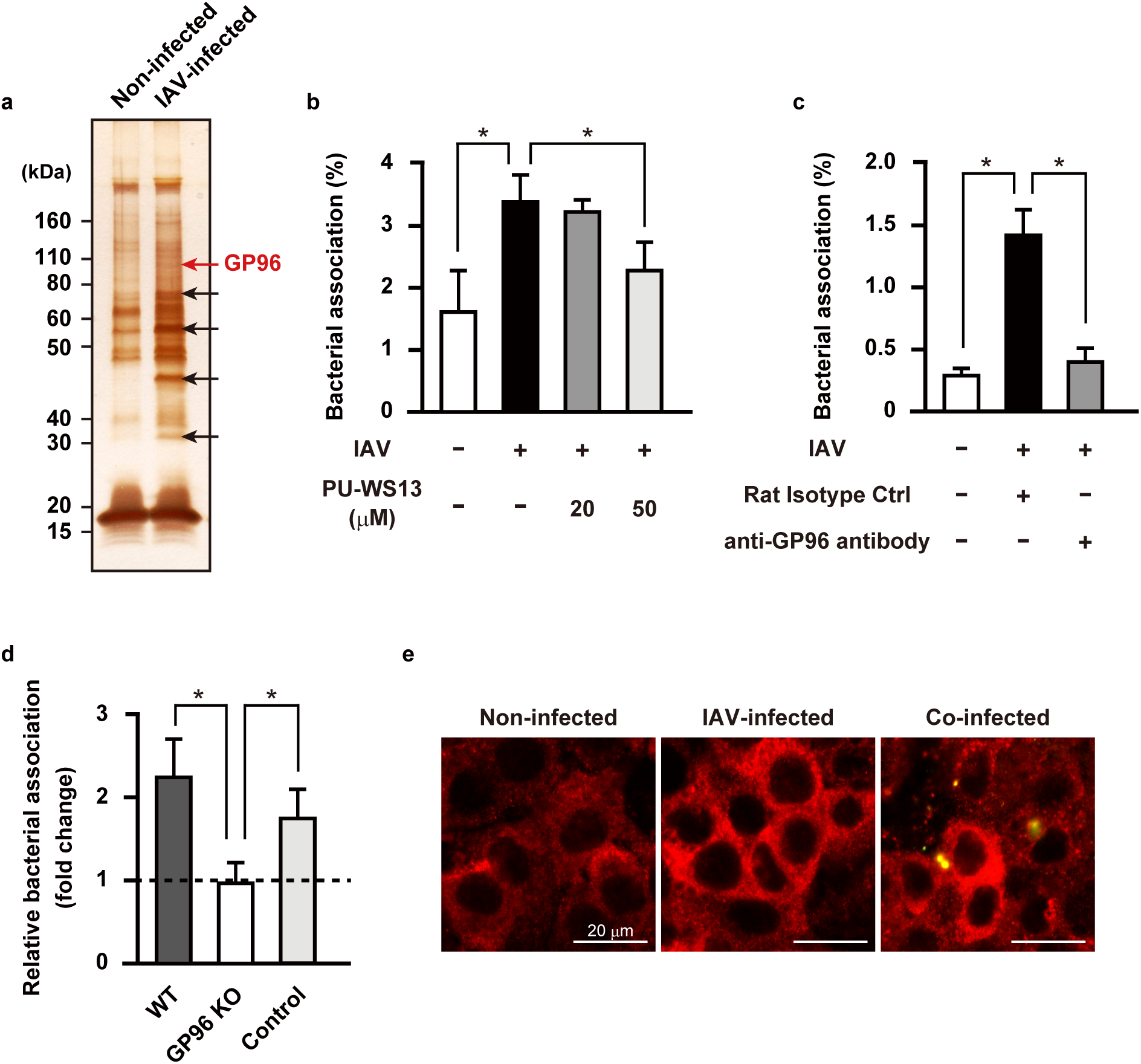
IAV infection-induced surface display of GP96 promotes pneumococcal adherence. **a**, A549 cells were infected with 10^6^ PFU of IAV for 36 hours, then treated with a membrane-impermeable biotinylation regent. Cell surface proteins were obtained using streptavidin beads, then subjected to SDS-PAGE and silver staining. Arrows indicate bands of upregulated surface proteins after IAV infection. **b, c**, A549 cells were infected with IAV for one hour. Following washing steps, they were incubated for 36 hours in the presence of PU-WS13 (**b**) or an antibody against GP96 (**c**). Next, IAV-infected cells were coinfected with *S*.*pneumoniae* D39 strain at an MOI of 5. At two hours after infection, cells were lysed and cell-associated bacteria were recovered. Bacterial adherence rate was calculated as percent of the inoculum. All experiments were performed in sextuplet with three technical repeats. Values are shown as the mean ± S.D. of six wells from a representative experiment. **d**, Effect of GP96 knockout on pneumococcal adherence. Bacterial association with IAV-infected cells was normalized to that with non-infected cells. All experiments were performed in sextuplet with three technical repeats. Values are shown as the mean ± S.D. of six wells from a representative experiment. \**P* <0.01 (**b**-**d**). **e**, A549 cells were infected with IAV followed by *S. pneumoniae* infection. GP96 was labeled with anti-GP96 and Alexa Fluor 594-conjugated antibodies (shown as red in images), while *S. pneumoniae* was labeled with anti-serotype 2 capsule and Alexa Fluor 488-conjugated antibodies (shown as green in images). Images were analyzed using a confocal laser scanning microscope. Values shown are representative of at least three separate experiments.

### Pneumococcal surface proteins AliA and AliB are determinants of bacterial adherence via GP96 receptor

In the early stage of infection, bacterial pathogens secrete a variety of virulence factors that interact with host receptors for establishment of colonization. To identify bacterial factors responsible for GP96-mediated adherence to IAV-infected epithelial cells, pneumococcal cell wall fractions were obtained and reacted with recombinant GP96 protein, then GP96-binding proteins were recovered by immunoprecipitation with an antibody against GP96. As shown in Figure 2a, protein bands with an apparent molecular mass of approximately 70 kDa were identified as AliA and AliB by mass spectrometry analysis. AliA and AliB are components of oligopeptide permease, and have functions related to bacterial colonization in the pharynx and lung^16^. To examine the interactions of each with GP96, immobilized recombinant Ali proteins and the predominant pneumococcal surface protein PhtD, used as a control, were incubated with serially diluted GP96, followed by detection with an anti-GP96 antibody. GP96 was found to bind to the AliA and AliB proteins, but not to PhtD (Fig. 2b). Furthermore, the binding affinity of Ali proteins to GP96 was evaluated using surface plasmon resonance (SPR) measurements. Equilibrium dissociation constants for the binding of AliA and AliB to GP96 protein were calculated by applying association and dissociation curves to a 1:1 Langmuir binding model (Table 1). SPR analysis revealed that both AliA and AliB bound to GP96 with a high affinity, with *K*_D_ values of 3.40 × 10^−8^ and 4.85 × 10^−8^ M, respectively. We next examined whether these pneumococcal Ali proteins function as adhesins for bacterial adherence to IAV-infected epithelial cells. Following IAV infection, the association of a wild-type (WT) strain to alveolar epithelial cells was increased by approximately 2.5-fold, whereas the adhesion activity of the *aliA* and *aliB* mutants remained unchanged (Fig. 2c). These results suggest that *S. pneumoniae* utilizes AliA and AliB proteins as adhesins to interact with surface-displayed GP96 on IAV-infected cells, resulting in establishment of a secondary pneumococcal infection.

**Table 1.** Kinetic binding parameters for pneumococcal surface protein to GP96

| Ligand | $k_a$ ($M^{-1} s^{-1}$) | $k_d$ ($s^{-1}$) | $K_D$ (M) |
| --- | --- | --- | --- |
| AliA | $2.31 \times 10^4$ | $7.85 \times 10^{-4}$ | $3.40 \times 10^{-8}$ |
| AliB | $2.09 \times 10^8$ | 10.12 | $4.85 \times 10^{-8}$ |

**Fig. 2.**
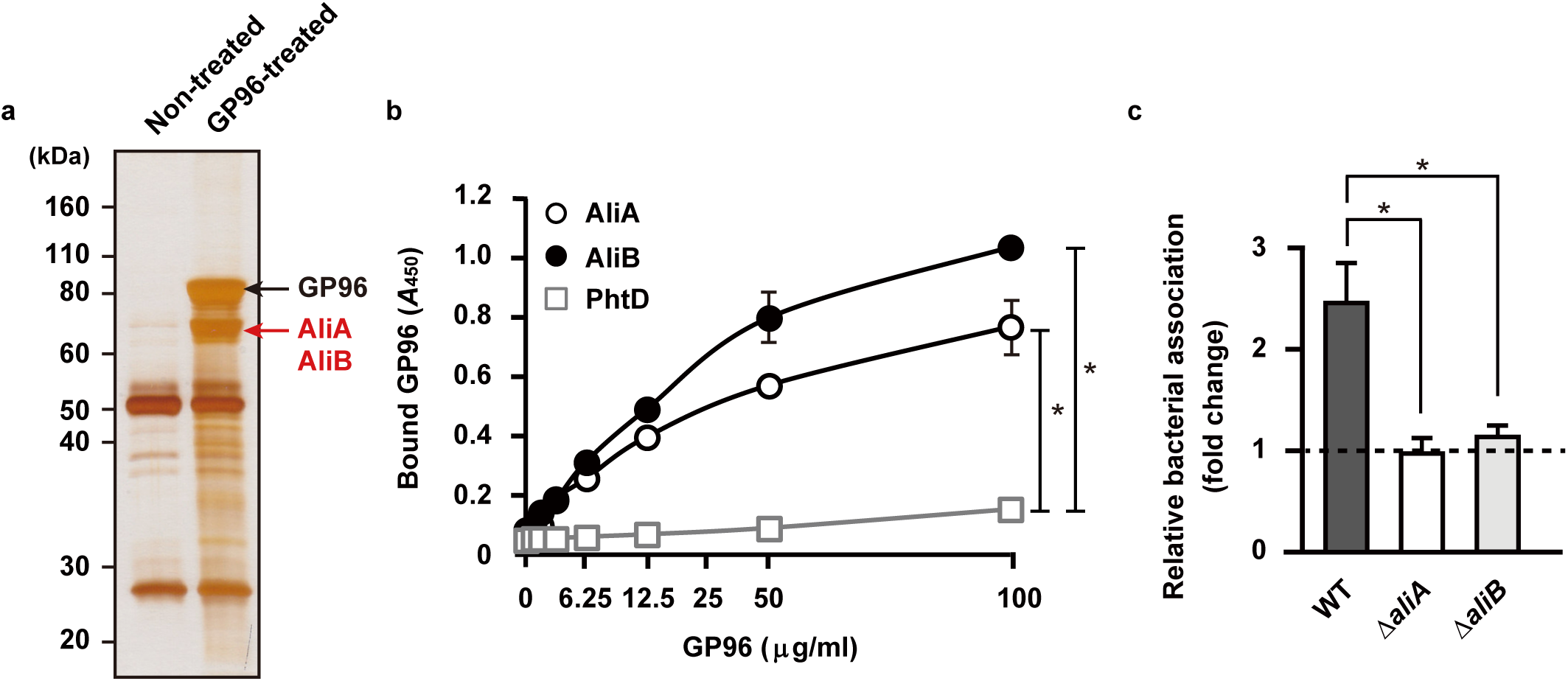
*S. pneumoniae* adheres to alveolar epithelial cells through interaction of pneumococcal surface proteins with GP96. **a**, Proteins bound to GP96 were immunoprecipitated from pneumococcal cell wall fractions, then subjected to SDS-PAGE and silver staining. **b**, AliA, AliB, and PhtD, bacterial surface proteins, were immobilized on microtiter plates, then increasing amounts of GP96 were added. Bound GP96 was detected using an anti-GP96 antibody. All experiments were performed in sextuplet with three technical repeats. Values are shown as the mean ± S.D. of six wells from a representative experiment. \**P* <0.01. **c**, Effects of deletion of *aliA* and *aliB* on pneumococcal adherence. Bacterial association with IAV-infected cells was normalized to that with non-infected cells. All experiments were performed in sextuplet with three technical repeats. Values are shown as the mean ± S.D. of six wells from a representative experiment. \**P* <0.01.

### Influenza infection-induced chaperoning activity of GP96 promotes pneumococcal adherence to alveolar epithelial cells

GP96 is a molecular chaperone that has a key role in folding as well as surface expression of various integrin subunits and Toll-like receptors (TLRs)^17^. Integrins are type I transmembrane heterodimeric proteins that mediate cell-cell and cell-extracellular matrix interactions. The major integrin ligand fibronectin (Fn) possesses a tripeptide arginine-glycine-aspartic acid (RGD) sequence that serves as the integrin recognition site. Bacterial pathogens responsible for secondary bacterial pneumonia, including *S. pneumoniae, Streptococcus pyogenes, Staphylococcus aureus*, and *Haemophilus influenzae*, utilize interactions with Fn-integrins to associate with host cells^18-21^. Thus, we speculated that IAV infection accelerates a GP96-dependent surface display of integrins that may augment bacterial association with epithelial cells. To examine in more detail, immunoprecipitation of cell lysates containing biotinylated surface proteins extracted from non-infected and IAV-infected epithelial cells was conducted using an antibody against integrin α_V_, a major α subunit involved in the pathogenesis of respiratory diseases. The expression levels of total integrin α_V_ were similar among all tested conditions, whereas a slight shift of the protein band potentially reflecting modification of sugar moiety was observed in IAV-infected cells (Fig. 3a, right panel). Notably, a marked increase in surface-exposed integrin α_V_ was seen following IAV infection, which was largely abrogated by introduction of the GP96 inhibitor, suggesting that integrin α_V_ is exported to the surface of IAV-infected cells in a GP96-dependent manner (Fig. 3a, left panel). Immunofluorescence staining experiments also revealed that co-localization of integrin α_V_ with GP96 on the cell surface was more pronounced with IAV infection (Fig. 3b).

**Fig. 3.**
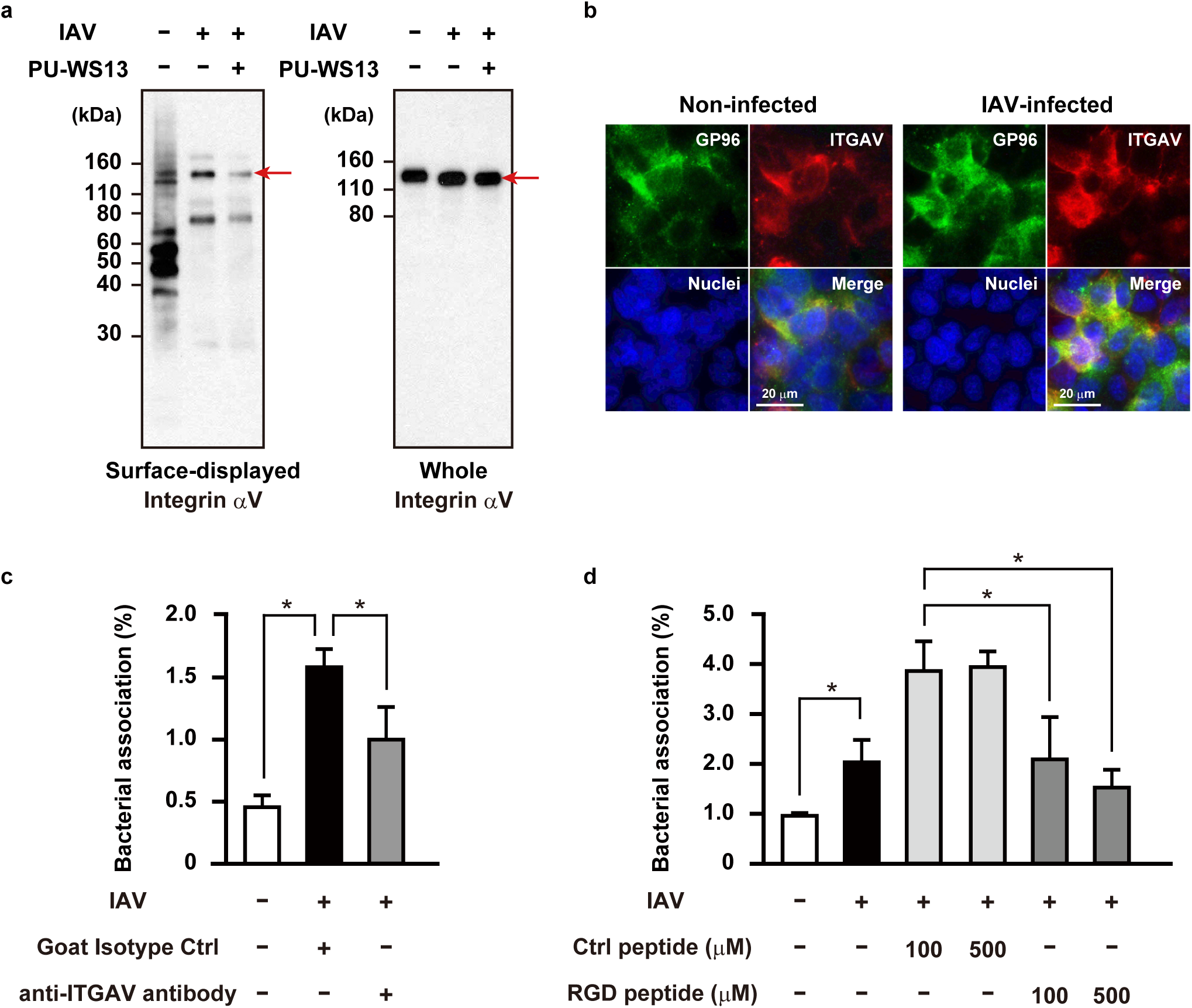
GP96-dependent surface display of integrin α_V_ associated with enhanced pneumococcal adherence following IAV infection. **a**, A549 cells were infected with IAV for 36 hours in the presence or absence of PU-WS13, then treated with a membrane-impermeable biotinylation reagent. Immunoprecipitation of cell lysates containing biotinylated surface proteins was performed using an antibody against integrin α_V_. Surface-displayed and whole-cell integrin α_V_ was detected using streptavidin and an antibody against integrin α_V_, respectively. Red arrows indicate integrin α_V_ band. **b**, A549 cells were infected with IAV for one hour. After transferring to fresh medium, incubation was continued for 36 hours. GP96 was labeled with anti-GP96 and Alexa Fluor 488-conjugated antibodies, while integrin α_V_ was labeled with anti-integrin α_V_ and Alexa Fluor 594-conjugated antibodies. DAPI was used to stain DNA in the nucleus. **c, d**, A549 cells were infected with IAV for 36 hours. Following washing steps, they were incubated with an antibody against integrin α_V_ (**c**) or RGD peptide (**d**) for one hour, then IAV-infected cells were coinfected with an *S*.*pneumoniae* strain at an MOI of 5. At two hours after initiating infection, cells were lysed and cell-associated bacteria recovered. Bacterial adherence rate was calculated as percent of inoculum. All experiments were performed in sextuplet with three technical repeats. Values are shown as the mean ± S.D. of six wells from a representative experiment. \**P* <0.01.

Next, we evaluated whether surface-exposed integrin α_V_ functions as a receptor for bacterial adherence to IAV-infected cells. As shown in Figure 3c, *S. pneumoniae* demonstrated a greater level of adherence to IAV-infected as compared to non-infected cells, though that enhanced bacterial association was partially reduced by treatment with an anti-integrin α_V_ antibody. Furthermore, as compared with a control peptide, an RGD-containing peptide abrogated pneumococcal adhesion in a dose-dependent manner (Fig. 3d). Thus, Fn-integrin interaction was also found to be associated with pneumococcal adherence to IAV-infected alveolar epithelial cells. We considered that the increased level of pneumococcal adherence to IAV-infected cells in the presence of control peptide was likely due to *S. pneumoniae* utilizing the synthetic peptides as a nutrient source. Together, these results provide evidence that IAV infection enhances the surface expression of integrin α_V_ through the chaperone activity of GP96, thereby increasing pneumococcal adherence to alveolar epithelial cells.

### Influenza infection-induced Snail1 expression contributes to disruption of alveolar epithelial barrier

Calcium (Ca^2+^) signaling has been implicated to be involved in various stages of host-pathogen interactions during viral and bacterial infections. Indeed, previous studies have shown that IAV infection induces Ca^2+^ influx, then elevated intracellular Ca^2+^ promotes endocytic uptake of the virus and host inflammatory response^22, 23^. Also, Ca^2+^ influxes activate calpains, Ca^2+^-dependent host cysteine proteases, which then target junctional proteins, such as occludin and E-cadherin^24^. Since calpains have been shown to change their distribution in an active state^25^, cellular localization of GP96 and calpains in IAV-infected alveolar epithelial cells was assessed using immunofluorescence staining. As opposed to GP96, calpain 2 was not only exposed on the apical surface of epithelial cells but also found concentrated in the plasma membrane region after IAV infection (Fig. 4a). Our observations suggest that IAV infection recruits calpains to the plasma membrane of paracellular junctions, which in turn promotes destabilization of the alveolar epithelial barrier through degradation of junctional proteins. We then evaluated the effects of IAV infection on expressions of GP96 and calpains, as well as junctional proteins associated with epithelial barrier function using quantitative RT-PCR analysis. Distinct downregulation of E-cadherin and p120-catenin was detected in IAV-infected cells, whereas that infection had no effect on expression of GP96 or calpains at the transcriptional level (Fig. 4b). These results imply that IAV infection does not induce increased expression levels of GP96 and calpains, but rather results in their ectopic localization in epithelial cells.

**Fig. 4.**
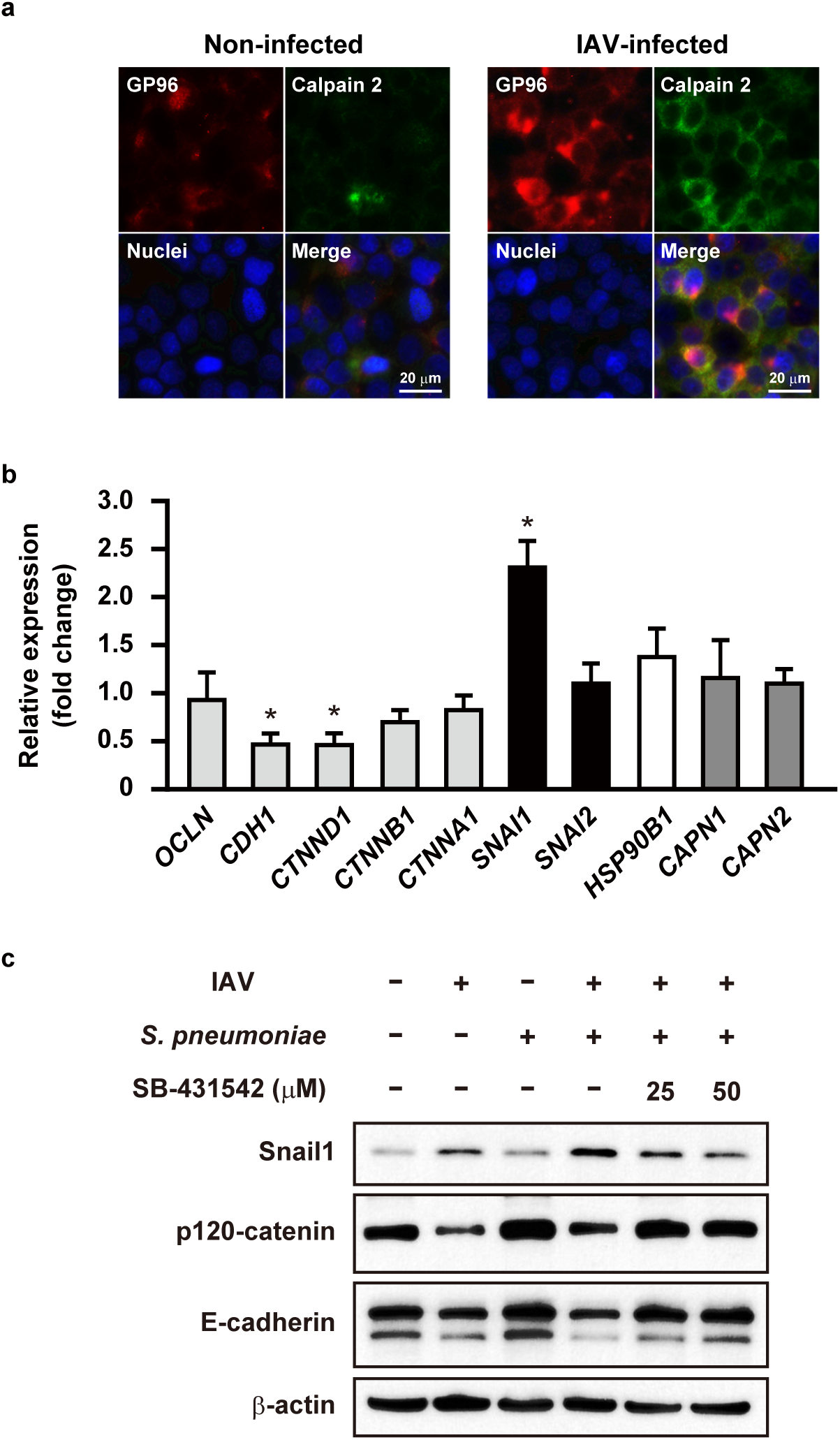
Calpain and Snail1 related to destruction of alveolar epithelial barrier following IAV infection. **a**, A549 cells were infected with IAV for 1 hour. After transferring to fresh medium, incubation was continued for 36 hours. GP96 was labeled with anti-GP96 and Alexa Fluor 594-conjugated antibodies, while calpains were labeled with anti-calpain 2 and Alexa Fluor 488-conjugated antibodies. DAPI was used to stain DNA in the nucleus. **b**, Transcriptional levels of genes encoding junctional proteins and regulators in A549 cells infected with IAV were analyzed using real-time RT-PCR. The *gapdh* transcript served as an internal control. Data from three independent tests are presented, with values shown as the mean ± SD for expression ratio. Transcriptional levels of tested genes are presented as relative expression levels normalized to that of non-infected cells. \**P* <0.01. **c**, A549 cells were infected with IAV for one hour, then incubated with fresh medium in the presence or absence of SB-431542. Following washing steps, cells were infected with an *S. pneumoniae* strain for seven hours. Expressions of E-cadherin, p120-catenin, and Snail1 were detected in whole cell lysates using Western blot analysis. β-actin served as a loading control.

Interestingly, IAV infection induced a drastic upregulation of the host transcriptional factor Snail1, a global repressor of genes encoding junctional proteins^26, 27^. TGF-β has been shown to downregulate the expression of E-cadherin via the snail signaling pathway, known to be fundamental for development of epithelial-to-mesenchymal transition (EMT)^28^. To investigate the association of Snail1 expression with destabilization of junctional proteins following infection, alveolar epithelial cells were infected with IAV in the presence or absence of a TGF-β inhibitor, then subsequently infected with *S. pneumoniae* (Fig. 4c). The protein level of Snail1 was elevated during both IAV infection and superinfection, whereas inhibition of the TGF-β pathway by SB-431542 in the co-infected cells resulted in remarkably reduced Snail1 levels in a dose-dependent manner. Along with increased levels of Snail1, reduced expression levels of E-cadherin and p120-catenin were observed in both IAV- and co-infected cells, while those were completely restored in the presence of the TGF-β inhibitor. Together, these data suggest that IAV infection induces Snail1-dependent dysfunction of the alveolar epithelial barrier via the TGF-β signaling pathway, thus providing a route for secondary pneumococcal dissemination.

### GP96 involved in exacerbation of bacterial pneumonia following influenza infection

To further clarify the role of IAV-induced GP96 in secondary pneumococcal pneumonia *in vivo*, mice were intranasally infected with a nonlethal dose of IAV (day 0), which was followed by intratracheal administration of the vehicle or GP96 inhibitor PU-WS13 on day 5, then intranasal challenge with *S. pneumoniae* was done on day 6 (Fig. 5a). At two days after bacterial infection, co-infected mice showed a significantly greater bacterial burden in the lungs as compared to those infected with *S. pneumoniae* only (Fig. 5b). Notably, PU-WS13 treatment resulted in a remarkable reduction in bacterial colonization in the lungs of the co-infected mice (Fig. 5b), indicating that mediation of GP96 induction by IAV infection has a key role in the pathogenesis of secondary bacterial pneumonia *in vivo*. Among integrin subsets, integrin α_V_β_6_ is an epithelium-restricted molecule expressed at low levels in the lungs of healthy individuals, then becomes rapidly upregulated in response to inflammation and injury. Meliopoulos, *et al*. reported that integrin β_6_ is an important factor associated with the severity of influenza diseases^29^. To further elucidate the mechanism by which IAV infection leads to increased susceptibility to secondary bacterial pneumonia, we examined the expression levels of integrin β_6_ and GP96 in pharyngeal and lung tissues at one day after bacterial administration under each infection condition (Fig. 5c, d). Quantitative RT-PCR analysis showed that IAV infection resulted in an approximately two-fold increase in expression levels of GP96 and integrin β_6_ in pharyngeal tissues (Fig. 5c), but not in lung tissues (Fig. 5d). Of note, high expression levels of GP96 and integrin β_6_ were detected in both pharyngeal and lung tissues after superinfection. These results suggest that pneumococcal colonization in the IAV-infected upper respiratory tract triggers GP96 expression in lung tissues, which in turn allows bacterial dissemination to the lower respiratory tract. Histopathological analysis was also performed using lungs obtained from mice at two days after pneumococcal infection (Fig. 5e). In lung tissues infected with IAV alone, moderate levels of inflammatory cell infiltration in peribronchiolar and interalveolar spaces, as well as microvascular hemorrhage were observed. Mice infected with *S. pneumoniae* alone also showed prominent perivascular and peribronchiolar lymphocytic cuffing. In contrast, mice with co-infection demonstrated dense leukocytic infiltration in interstitial and alveolar spaces, along with hemorrhaging, vascular leakage, and edema formation, suggesting vascular damage and increased epithelial-endothelial permeability. Notably, PU-WS13 treatment resulted in improvement of lung pathology in co-infected mice. To investigate lung tissue integrity in superinfected mice, we examined the expression of E-cadherin in lung tissues obtained from IAV-as well as co-infected mice. Those with co-infection exhibited a marked decrease in E-cadherin expression as compared to mice infected with IAV alone, while that was partially restored by PU-WS13 treatment (Fig. 5f). Taken together, our results indicate that GP96 is a factor for exacerbation of secondary bacterial pneumonia following influenza as well as a promising novel target for therapeutic intervention.

**Fig. 5.**
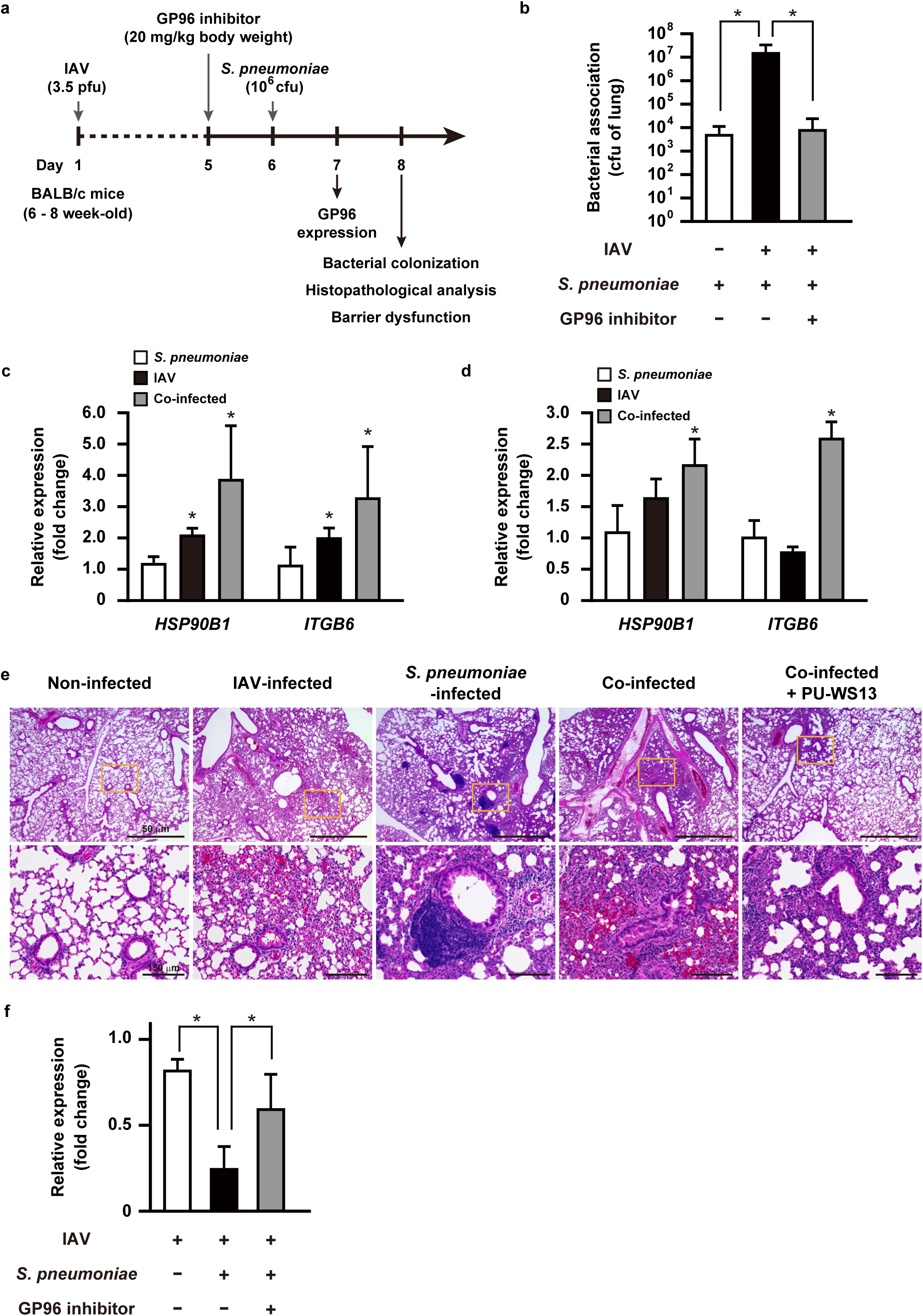
GP96 a crucial factor for exacerbation of bacterial pneumonia following IAV infection. **a**, Schematic showing experimental design. Mice were intranasally infected with IAV (day 0), followed by *S. pneumoniae* on day 6. In some experiments, PU-WS13, a GP96 inhibitor, was intratracheally administered. **b**, Effect of PU-WS13 treatment on bacterial burden in lungs. Values shown represent the mean ± S.D. of quintuplet samples and are representative of at least three independent experiments. \**P* <0.01. **c, d**, Transcriptional levels of genes encoding GP96 and integrin β_6_ in pharyngeal (**c**) and lung tissues (**d**) infected with IAV- and *S. pneumoniae* were analyzed by real-time RT-PCR. The *gapdh* transcript served as an internal control. Values from three independent tests are presented as the mean ± SD for expression ratio. \**P* <0.01. **e**, Lung tissues obtained from mice infected under various conditions were subjected to hematoxylin and eosin staining. Boxed area is magnified and shown in lower panel. **f**, Transcriptional levels of E-cadherin gene in lung tissues infected under various conditions were analyzed by real-time RT-PCR. The *gapdh* transcript served as an internal control. Values from three independent tests are shown as the mean ± SD for expression ratio. Transcriptional levels are presented as relative expression levels normalized to that of non-infected tissues (**c, d, f**). \**P* <0.01.

## Discussion

Secondary bacterial infections following a primary influenza virus infection are frequent complications, and result in the majority of related deaths during seasonal and pandemic influenza outbreaks. *S. pneumoniae* is the most commonly identified pathogen in secondary bacterial pneumonia cases. Although antibiotics remain the mainstay of therapy for affected patients, the increasing prevalence of multidrug-resistant *S. pneumoniae* is a serious public health concern worldwide. Thus, development of host-directed therapeutics is receiving focus as an alternative approach to treating secondary bacterial pneumonia following influenza. The present findings showed that GP96 functions as an exacerbation factor for secondary bacterial infections following influenza, thus we propose GP96 as a potential therapeutic target for novel countermeasures used to treat bacterial pneumonia.

GP96, an ER-resident HSP90 paralogue, has been reported to be exposed on the surface of various types of cells by multiple types of microbial infection^13^. The present is the first study to show that IAV infection triggers surface distribution of GP96 in human airway epithelial cells, where it is then hijacked as a host receptor for secondary infection by *S. pneumoniae*. The interaction between extracellular GP96 and bacterial surface ligands has been shown to activate host signaling cascades that facilitate bacterial adherence and internalization. Indeed, pathogenic *Escherichia coli* targets Ecgp96, a homologue of GP96 expressed on human brain microvascular endothelial cells, induces rearrangement of actin microfilaments and disassembly of endothelial junctions through signaling-mediated Ca^2+^ influx and nitric oxide production, resulting in acceleration of bacterial invasion^30, 31^. Bacterial pore-forming toxins such as pneumolysin produced by *S. pneumoniae* also induce similar cellular events in host cells^32^. Although the present results demonstrated that IAV infection induces Ca^2+^-dependent calpain activation, which then evokes destabilization of paracellular junctions without bacterial infection, *S. pneumoniae* may utilize not only extracellular GP96-mediated signaling but also pneumolysin-induced cell damage for invasion into deeper tissues. Despite increased understanding regarding the roles of GP96 in the pathogenesis of infections, the molecular mechanisms underlying surface distribution remain unidentified. Recently, plasma membrane damage mediated by bacterial pore-forming toxins was shown to be associated with redistribution of GP96 via non-muscle myosin II activity and Ca^2+^ influx during *Listeria monocytogenes* infection^33^. Although the mechanism governing the surface distribution of GP96 following IAV infection remains largely unknown, it is likely that calcium homeostasis and Ca^2+^-dependent effectors have key roles in IAV-infected airway epithelial cells. Bacterial colonization in the upper respiratory tract is considered to be a prerequisite for invasive infection, which results in bacterial invasion into other tissues or dissemination to the lower respiratory tract. In addition to a drastic increase in receptor availability accompanied by influenza virus infection, *S. pneumoniae* also possesses a variety of adhesins that augment bacterial adherence to these newly exposed receptors^34^. Herein, we identified AliA and AliB as bacterial adhesins for the display of GP96 on the surface of alveolar epithelial cells following IAV infection. These oligopeptide-binding proteins are conserved among bacterial pathogens most frequently associated with influenza, including *S. pneumoniae, S. pyogenes, S. aureus*, and *H. influenza*. GP96 serves as the host cellular receptor for various bacterial adhesins, such as pathogenic *E. coli* OmpA^30, 31^, *L. monocytogenes* Vip^14^, *S. aureus* Bap^35^, and *Clostridium difficile* enterotoxin A^36^, though those proteins share no homology with Ali proteins. Since GP96 might nonspecifically interact with multiple bacterial molecules on account of its chaperone structure, bacterial pathogens likely utilize ectopically exposed GP96 for establishment of bacterial colonization.

Transforming growth factor (TGF-β), a multifunctional cytokine, is secreted in an inactive or latent form, then subsequently activated through various mechanisms. During IAV infection, viral neuraminidase was shown to activate TGF-β, which promoted upregulation of host adhesion molecules, including fibronectin and integrins^37^. TGF-β is also a positive regulator of the integrin signaling pathway that promotes cytoskeletal rearrangement and bacterial internalization^38^. Indeed, we previously reported that *S. pyogenes* possesses Fn-binding molecules and utilizes Fn-integrin interactions to adhere to and invade IAV-infected cells^39^. In addition to TGF-β signaling, the present findings showed that IAV infection induces upregulation and display of integrins on the surface of alveolar epithelial cells via a chaperoning activity of GP96. Furthermore, GP96 serves as an essential chaperon for the cell-surface protein glycoprotein A repetitions predominant (GARP), a docking receptor for latent membrane-associated TGF-β^40^, indicating GP96 as a crucial factor for TGF-β signaling. Notably, *S. pneumoniae* also expresses neuraminidases on bacterial cell walls. NanA, a primary pneumococcal neuraminidase, is a sialidase that catalyzes cleavage of terminal sialic acids from latency-associated peptide (LAP) and also activates TGF-β signaling^41^. Another study showed that activation of TGF-β signaling proceeds through phosphorylation of SMAD proteins, which is associated with Snail1-mediated down-regulation of tight junction proteins of epithelial and endothelial cells^42^. The present results provide evidence that a preceding influenza infection induces a Snail1-dependent dysfunction of the airway epithelial barrier through TGF-β signaling, thus preparing a route for secondary pneumococcal translocation into deeper tissues via paracellular junctions. Therefore, the synergistic effects of the GP96 chaperoning function and viral-pneumococcal neuraminidase activities likely prime potent TGF-β signaling, which leads to increased bacterial loading in the lungs and pulmonary barrier dysfunction.

Besides being exploited as a host receptor for variety of pathogens, GP96 modulates host immune response to counteract an infection. GP96 functions as a master chaperone for cellular localization and function of TLRs. During an influenza infection, TLRs act as key transducers of type I interferons (IFNs) by recognition of viral nucleic acid^4^. It is known that type I IFN signaling through the IFN-α/β receptor evokes expression of proinflammatory genes to inhibit viral replication and augment various aspects of adaptive immunity, while several lines of evidence also indicate that neutrophil function is impaired following influenza virus infection. Viral infection-primed expression of type I IFNs is sufficient to interfere with production of specific chemokines, such as CXCL1 and CXCL2, resulting in impairment of neutrophil response during secondary *S. pneumoniae* infection^43^. The present findings showed that pneumococcal colonization in the upper respiratory tract triggered upregulation of GP96 in murine pharyngeal and lung tissues infected with IAV through an unidentified mechanism. Accordingly, given the importance of GP96 in TLRs signaling^17^, IAV infection-induced GP96 might impair immune responses against *S. pneumoniae* by excess production of type I IFNs. On the other hand, GP96 has been shown to specifically bind to and activate neutrophils and monocytes^44^. Nevertheless, we speculate that binding of pneumococcal Ali proteins to GP96 interferes with direct interactions of GP96 with neutrophils as well as monocytes in cases of viral-bacterial dual infection. Although direct evidence showing that GP96 is related to impairment in macrophages and neutrophil responses following IAV infection remains lacking, it may function as a potent receptor for bacterial colonization as well as an immune regulator in dysfunction of innate immune defenses against bacterial infection.

Taken together, the present results indicate that GP96 functions as a multifunctional exacerbation factor to promote pneumococcal colonization dysfunction of lung tissue, barrier, and probably dysregulation of immune responses as well, during secondary bacterial pneumonia following an influenza infection. Because of the complexity of the pathogenesis, a balanced combination of antimicrobial agents and immunomodulators could be more effective for prospective therapeutics. We believe that GP96 is a potential target for development of promising therapeutic strategies, including combination therapies as alternatives to conventional antibiotics and antiviral agents administered for broad-spectrum prevention, as well as management of secondary bacterial infections following influenza.

## Methods

### Bacterial and viral strains and culture conditions

*Streptococcus pneumoniae* D39 (serotype 2 clinical isolate) and isogenic mutant strains were cultured in Todd-Hewitt broth (Becton, Dickinson and Company; BD) supplemented with 0.2% yeast extract (BD) (THY medium) at 37°C in an ambient atmosphere. For selection and maintenance of mutant strains, spectinomycin (Sigma-Aldrich) at 100 μg/ml was added to the medium. *Escherichia coli* strains BL21-gold (DE3) (Agilent Technologies) and XL10-gold (Stratagene) served as hosts for derivatives of plasmids pGEX-6P-1 (Cytiva) and pQE30 (Qiagen), respectively. All *E. coli* strains were cultured in Luria-Bertani (Nacalai Tesque) (LB) medium at 37°C with agitation. For selection and maintenance of *E. coli* mutant strains, ampicillin (100 µg/ml) was added to the medium. Influenza A virus A/FM/1/47 (H1N1) was grown in Madin-Darby canine kidney (MDCK) cells.

### Cell cultures and construction of GP96 knockout cells

The human alveolar carcinoma cell line A549 (Riken Cell Bank) derived from type II pneumocytes and MDCK were maintained in DMEM supplemented with 10% FBS at 37°C under a 5% CO_2_ atmosphere.

A CRISPR-Cas9 GP96 knockout plasmid was created by cloning sgRNA GP96 oligos into a pSpCas9(BB)-2A-Puro (PX459) plasmid, a gift from Feng Zhang (Addgene plasmid #62988)^45^. The constructed plasmid was then transfected into A549 cells with Lipofectamine 3000 Reagent (Thermo Scientific), according to the manufacturer’s instructions. At 24 hours after transfection, 1.5 µg/ml puromycin was added and the cells were further cultured for two days. Next, a limiting dilution of the survived cells was conducted and single colonies were expanded, with GP96 expression examined by Western blot analysis using anti-Grp94 rabbit Ab (Cell Signalling). PCR products were amplified with purified genomic DNA, and the primers gp96checkF and gp96checkR, then subjected to Sanger sequencing for confirmation of mutations. All primers used are listed in Supplementary Table 1.

### Preparation of *S. pneumoniae* mutant strains and recombinant proteins

Inactivation of the *aliA* and *aliB* genes was achieved by transformation of strain D39 with a linear DNA fragment containing a spectinomycin resistance gene (*aad9*) flanked by the upstream and downstream sequences of the *aliA* and *aliB* genes, as previously reported^46^.

For construction of recombinant GP96, cDNA of A549 cells was prepared using Trizol and a PureLink RNA mini-kit (Thermo Scientific). cDNA fragments encoding full-length GP96 were amplified using specific primers (Supplementary Table 1). The fragments were cloned into a pGEX-6P-1 vector via the *BamH*I and *Sal*I sites, then transformed into *E. coli* BL21 (DE3).

Recombinant AliA and AliB proteins were hyper-expressed in *E. coli* XL10-Gold using a pQE30 vector. N-terminal His-tagged proteins were purified using a QIAexpress protein purification system (Qiagen), as previously described^47^.

### Adherence assay

A549 cells were cultured in 24-well plates at a density of 2 × 10^5^ cells per well and infected with 10^6^ PFU of IAV in serum-free DMEM supplemented with 0.1% BSA (Sigma-Aldrich), MEM vitamin solution (Thermo Scientific), and 0.01% DEAE dextran (GIBCO) for one hour at 34°C. Following washing steps, the cells were incubated for 36 hours in the presence or absence of PU-WS13 (Merck Millipore). *S. pneumoniae* strains were grown to the exponential phase (OD_600_ = 0.7), then washed with and resuspended in PBS. IAV-infected cells were exposed to 10^7^ CFU of pneumococci in DMEM supplemented with 10% FBS for two hours at 37°C. For quantification of bacterial adherence, infected cells were washed with PBS and lysed with distilled water. Serial dilutions of the lysates were plated on THY agar plates to determine CFU.

In some experiments, antibodies targeting Grp94 (Rat, mAb, Enzo) and integrin α_V_ (Goat, pAb, R&D), rat IgG2A isotype control (Rat, mAb, R&D), normal goat IgG control (Goat, pAb, Enzo), RGD peptide (Gly-Arg-Gly-Asp-Asn-Pro, Enzo), and RGD control peptide (Gly-Arg-Ala-Asp-Ser-Pro, Enzo) were added to A549 cells at one hour before infection with *S. pneumoniae*.

### Immunoprecipitation assay

Cell surface proteins were biotinylated using an EZ-Link Sulfo-NHS-Biotin reagent (Thermo Scientific) following the manufacturer’s protocol. Briefly, non-infected and IAV-infected A549 cells were washed with PBS, then incubated with 1 mM Sulfo-NHS-biotin for 30 minutes at room temperature. After quenching the biotin reagent with 100 mM glycine in PBS, the cells were lysed using radioimmunoprecipitation buffer containing Complete protease inhibitor (Roche Life Science) for 20 minutes at 4°C. Cell surface proteins were precipitated with Dynabeads M-280 Streptavidin (Thermo Scientific), separated using SDS-PAGE, then visualized by silver staining. Bands of interest were analyzed by liquid chromatography-tandem mass spectrometry using a Q-Exactive Mass Spectrometer (Thermo Scientific) equipped with an UltiMate 3000 Nano LC System (Thermo Scientific). Raw data were processed using Mascot Distiller v2.5 (Matrix Science). Peptide and protein identification was performed with Mascot, v.2.5 (Matrix Science), using the UniProt database with a precursor mass tolerance of 10 ppm, a fragment ion mass tolerance of 0.01 Da, and strict trypsin specificity that allowed one missed cleavage. To determine cellular localization of integrin α_V_, immunoprecipitation using an antibody against integrin α_V_ (Rabbit, pAb, Cell signaling) and Dynabeads Protein A (Thermo Scientific) was performed, then detection was done by Western blot analysis with a specific antibody against integrin α_V_ (Goat, pAb, R&D) and a horseradish peroxidase (HRP)-conjugated antibody against goat IgG (R&D), or HRP-conjugated streptavidin (Thermo Scientific). Immunoreactive bands were detected using Pierce Western blotting substrate (Thermo Scientific).

For identification of bacterial factors associated with GP96, cell wall fractions of *S. pneumoniae* were prepared as previously described^49^. The fractions were incubated with 10 μg of recombinant GP96 for six hours at 4°C. Proteins bound to GP96 were immunoprecipitated with an antibody against Grp94 (Rat, mAb, Enzo) and Dynabeads Protein G (Thermo Scientific), then examined using mass spectrometry analysis, as described above.

### Surface plasmon resonance analysis

Association and dissociation reactions of GP96 to pneumococcal Ali proteins were analyzed using a BIAcore optical biosensor (BIAcore X-100 system, GE Healthcare Life Sciences), as previously described^48^. Briefly, recombinant GP96 (20 μg ml^−1^ in 10 mM sodium acetate, pH 4) was covalently immobilized on a CM5 sensor chip using an Amine coupling kit (GE Healthcare Life Sciences). Binding analyses were performed in HBS-P buffer (0.01 M HEPES, pH 7.4, 0.15 M NaCl, 0.005% surfactant P20; GE Healthcare Life Sciences) at 37°C with a flow rate of 30 μl/min. AliA and AliB were separately used as an analyte at concentrations of 31.3, 62.5, 125, 250, and 500 nM. Parameters of binding kinetics were analyzed using raw data from the BIAcore sensorgram suitable for analysis with the kinetic models included in the BIA evaluation software package, v. 3.0.2 (GE Healthcare Life Sciences). Data were fitted using a 1:1 Langmuir binding model.

### ELISA

GP96 binding to AliA and AliB proteins was assessed by ELISA, as previously described^50^. Microtiter plates (96-well; Sumitomo Bakelite) were separately coated with AliA, AliB, and PhtD protein (250 ng) in coating buffer (0.1 M Na_2_CO_3_, 0.1 M NaHCO_3_, pH 9.6) at 4°C overnight. The plates were blocked with 10% Block Ace solution (Megmilk Snow Brand) at 4°C overnight, then washed with PBS containing 0.2% Tween 20 (PBST). GP96 was diluted with binding buffer (50 mM HEPES, pH 7.4, 150 mM NaCl, 2 mM CaCl_2_, 50 μg/ml BSA) and incubated with immobilized bacterial surface proteins for 90 minutes at 37°C. After washing with PBST, the plates were incubated with an antibody against GP96 (Sheep, pAb, R&D) for two hours at room temperature. Subsequently, an HRP-conjugated antibody against sheep IgG (R&D) was added to the plate and incubation was continued for two hours at room temperature. Following a washing step, the peroxidase substrate tetramethylbenzidine (Moss) was added to the plate. The reaction was stopped by addition of 0.5 N HCl and absorbance at 450 nm was measured using a Muktiskan FC microplate photometer (Thermo Scientific).

### Immunofluorescence microscopy

A549 cells were seeded at 2 × 10^5^ onto cover slides (13-mm diameter; Matsunami) pretreated with coating buffer containing 0.1% collagen (Type I from rat tail; Sigma-Aldrich) and 0.1% gelatin (from bovine bone; Wako). The cells were then infected with IAV followed by *S. pneumoniae*, as described above, and fixed with 4% paraformaldehyde-PBS. Following blocking with 5% bovine serum albumin-PBS, the cells were reacted with a primary antibody targeting Grp94 (Rat, mAb, Enzo), Grp94 (Rabbit, pAb, Thermo Scientific), integrin α_V_ (Mouse, mAb, R&D), Calpain 2 (Rabbit-pAb, Cell Signaling), or serotype 2 capsule (Rabbit, pAb, Denka Seiken). After washing steps, incubation was performed with Alexa Fluor 594-conjugated anti-rat IgG (Thermo Scientific), Alexa Fluor 594-conjugated anti-mouse IgG (Thermo Scientific), Alexa Fluor 488-conjugated anti-mouse IgG (Thermo Scientific), or Alexa Fluor 488-conjugated anti-rabbit IgG (Thermo Scientific). To observe the association of pneumococci with GP96, imaging was performed using a Zeiss LSM 510 confocal microscope system, v. 3.2 (Carl Zeiss) and analyzed with the LSM 510 software package. For assessment of cellular localization of GP96, integrin α_V_, and calpain 2, cover slides were enclosed with ProLong Gold Antifade Reagent with DAPI (Thermo Scientific) and examined using a Carl Zeiss Axioplan 2 fluorescent microscope system.

### Analysis of destabilization of epithelial junctions

Whole cell lysates from coinfected epithelial cells were prepared as previously described^49^. Briefly, A549 cells were infected with IAV in the presence or absence of SB-431542, a TGF-β inhibitor. At seven hours after infection, the cells were lysed with Laemmli gel loading buffer containing 6% 2-mercaptoethanol. Cleavage of junctional proteins was detected by Western blot analysis using specific antibodies against E-cadherin (Mouse, mAb, Thermo Scientific), p120-catenin (Rabbit, pAb, Cell Signaling), Snail1 (Mouse, mAb, Cell Signaling), and β-actin (rabbit, pAb, Cell Signaling). Horseradish peroxidase (HRP)-conjugated antibodies against mouse or rabbit IgG (Cell Signaling) were used as the secondary antibodies.

### Mouse experiments

Female BALB/c mice at six to eight weeks old (CLEA Japan, Inc.) were intranasally infected with 3.5 PFU of IAV A/FM/1/47 (H1N1) in 40 μl PBS (day 0). The *S. pneumoniae* D39 strain was grown to the mid-exponential phase (OD_600_ = 0.4), then washed with and resuspended in PBS. Bacteria were introduced into the mice by intranasal administration of 1 × 10^6^ CFU in 40 μl PBS on day six. In some experiments, mice were intratracheally treated with either the vehicle or PU-WS13 (20 mg/kg mouse body weight) on day five.

For quantification of bacterial colonization in the lung, mice were euthanized two days after *S. pneumoniae* infection, then lung tissues were immediately collected. Lung homogenates were serially diluted and plated on THY agar plates containing 5% sheep blood. For histopathologic examinations, lung tissue samples were obtained and fixed with formalin, then embedded in paraffin and sectioned, and subjected to hematoxylin and eosin (HE) staining. Stained tissue sections were observed using an EVOS M5000 cell Imaging system (Thermo Scientific). For assessment of gene expression, pharyngeal and lung tissues were harvested at various time points.

### Real-time RT-PCR assay

Total RNA was isolated from A549 cells using a CellAmp Direct RNA Prep Kit (TaKaRa), as well as murine pharyngeal and lung tissues using an RNeasy Fibrous Tissue Mini Kit (QIAGEN). Synthesis of cDNA from total RNA was performed with a PrimeScript RT reagent Kit (TaKaRa). The possibility of DNA contamination was excluded by PCR analysis of non-RT samples. Primer sets for selected genes were designed using Primer Express Software, version. 3.0 (Applied Biosystems). All primers used are listed in Supplementary Table 1. RT-PCR amplifications were performed using the SYBR Green method with an ABI StepOne™ Real-Time PCR System, v. 2.2 (Applied Biosystems). Relative expression amounts were calculated with the ΔΔC_T_ method. The level of *gapdh* expression was used as an internal control.

### Statistical analysis

All statistical analyses were performed using GraphPad Prism, v. 7.03 (GraphPad Software). Differences were determined with Mann-Whitney’s *U* test when comparing two groups, or one-way ANOVA, followed by Tukey’s multiple comparison test when comparing multiple groups. A confidence interval with a *P* value of <0.01 was considered to be significant.

### Ethics statement

All mouse experiments were conducted using a protocol approved by the Animal Care and Use Committee of Osaka University Graduate School of Dentistry (authorization number 01-010-0) and the Animal Care and Use Committee of Kanazawa University (authorization numbers AP-143262 and AP-183936).

## Acknowledgements

This work was supported by AMED (grant number JP19fm0208007), and grants from JSPS KAKENHI (JP19H03825, JP18K19643, JP17K11610, JP17H04369, JP17K19751, JP17H05103, JP18K17027), as well as a GSK Japan Research Grant and the Takeda Science Foundation, Kobayashi International Scholarship Foundation, and Naito Foundation. We thank A. Bando for the helpful technical assistance.

## Authors’ contributions

T.S. and S.K. conceived and designed the experiments. T.S., M.N., S. N., Y. T., M.H.O., and Y.M. performed the experiments. T.S., M.N., M.Y., and S.O. analyzed the data. T.S. and M.N. contributed to writing of the manuscript. All authors participated in discussions related to the present research, and reviewed and approved the final version of the manuscript.

## Sumitomo et al. Supplemental table 1

**Table S1.**
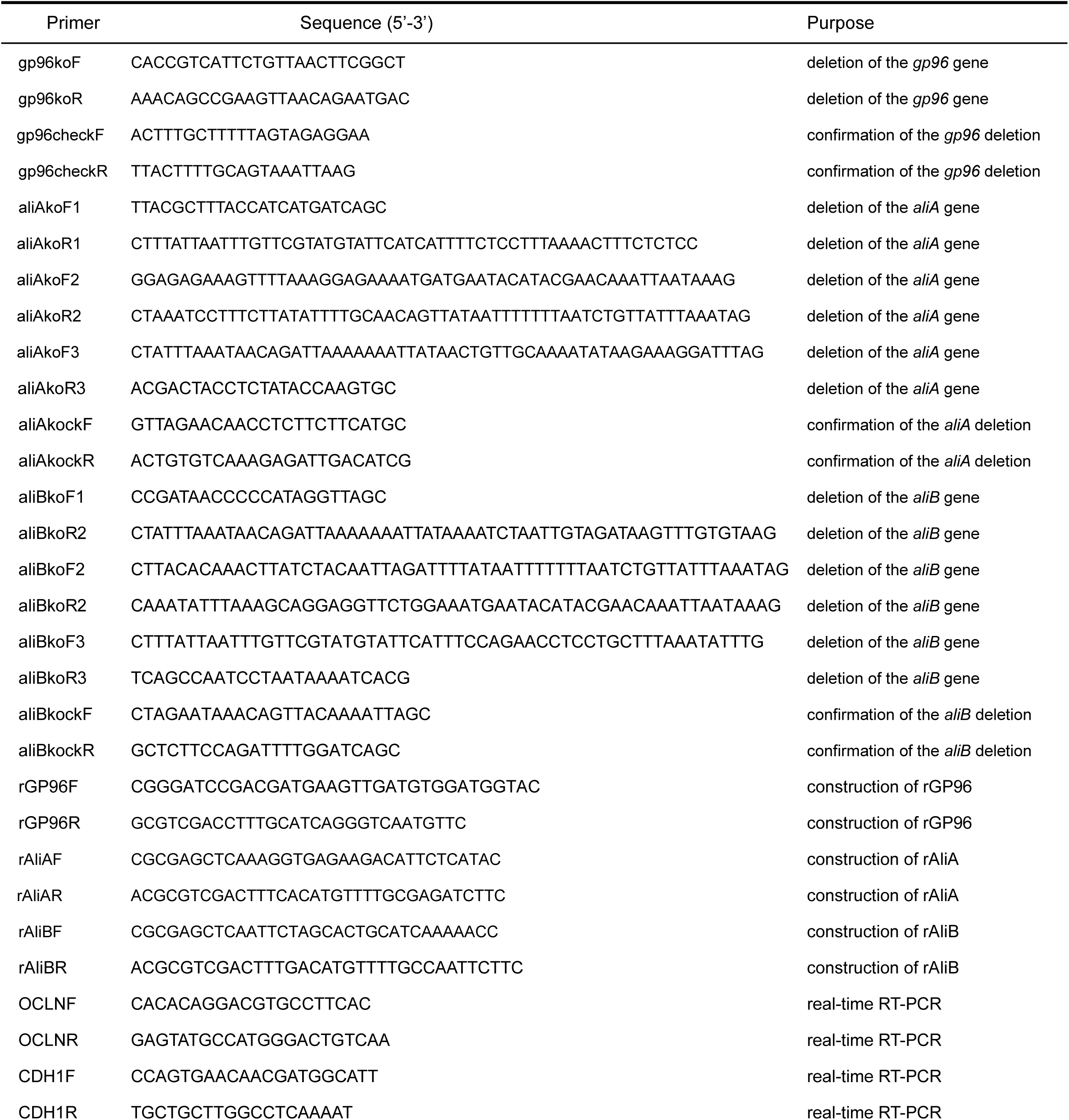
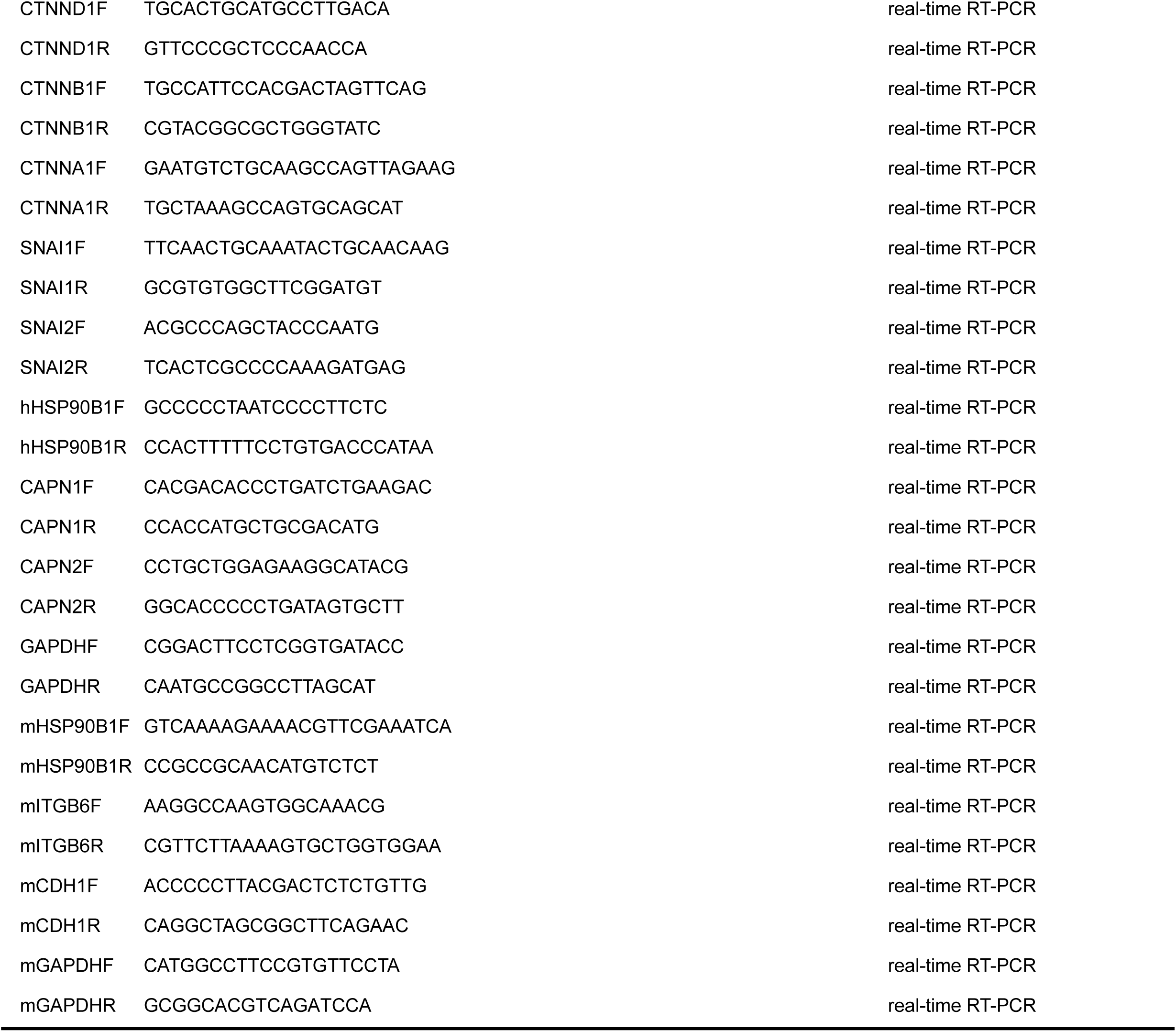
Oligonucleotides used in this study.

